# Locating Evolutionary Rate Inflection Points on Whole-Genome SNV Similarity Curves and Their Application in Identifying Key Mutations in Language/Cognition Genes

**DOI:** 10.64898/2026.08.05.743118

**Authors:** Zhizhou Zhang, Yongdong Xu

## Abstract

Language genes can be tentatively considered as a subset of cognitive genes, although they are often discussed separately. During the evolution of SNVs (single nucleotide variations) in cognition-related genes, do language genes and cognitive genes exhibit significantly different intensities of change at several key evolutionary moments—namely, the inflection points or derivative peak positions of similarity curves drawn from multi-SNV locus bases across samples? In this study, nine distance/similarity metrics (Bray–Curtis, Cosine, Pearson, Spearman, Hamming, Jaccard, Matching, Kulczynski, and Gower) were employed to analyze 413 samples from 11 taxonomic groups, targeting SNV loci in language/cognition-related genes (13,415 effective loci, approximately 400 loci per gene), with pp6 (Homo_sapiens.GRCh38) as the reference. For each method, sample similarities (defined as 1/(1+distance)) were independently sorted in ascending order to generate raw similarity scatterplots. Due to the large sample size and representativeness, the scatter density on the similarity curves was high, and no smoothing was applied. Derivative values were calculated from adjacent similarity differences to identify peaks of evolutionary rate change (top 10 peaks per method). Combined with functional annotations of 33 language/cognition-related genes, we quantified the difference scores and occurrence frequencies of the two gene categories at the peak positions. The results indicate that cognitive-related genes exhibit slightly higher occurrence frequencies in peak windows and higher average difference scores per gene than language genes. Comparative analysis of SNVs at the peak samples and their left-side windows revealed that at positions 381–382, all nine methods shared three intersecting mutation loci, involving language genes (NFXL1, SRGAP2, SRGAP2C); at positions 355–356, there was one intersecting mutation locus, involving a language gene (SRGAP2). This suggests that certain mutations in language genes may have played a distinctive role at critical junctures in the evolution of cognitive abilities.

## Introduction

The relationship between language and cognition has become increasingly important in the current era of artificial intelligence [1–2]. When placed in the broad evolutionary perspective from fish to modern humans, a picture emerges in which continuity and uniqueness coexist: the neural and genetic foundations of basic language abilities are extremely ancient, with homologs found even in distantly related animals such as fish. Foundational cognitive capacities like category learning are widespread across the animal kingdom and constitute the cognitive bedrock for language evolution. Meanwhile, various components of higher-level language abilities—such as referentiality and syntactic rules—are already evident to varying degrees in birds and mammals. However, the integration of these components into a uniquely human symbolic system with unlimited combinatorial capacity likely represents a leap achieved during the later stages of human evolution. Thus, from fish to modern humans, language ability is not an all-or-nothing mutation but rather a long story of an ancient biological foundation being continuously modified, enriched, and reorganized [3–5]. A similar evolutionary blueprint appears to apply to the evolution of cognitive abilities as well. Cognitive abilities include basic cognition (e.g., sensation, perception, attention) and higher-order cognition (e.g., reasoning, association, complex mathematical calculation); the neural and genetic foundations of basic cognition are also extremely ancient and highly conserved from fish to humans. The core mechanisms of abilities such as perception, learning, and memory have maintained remarkable continuity over hundreds of millions of years of evolution. Components of higher-order cognition—such as tool use and social cognition—are already manifested to varying degrees in birds and fish. Yet the integration of these components into a uniquely human cognitive system with unlimited creativity and abstract thinking capacity also represents a major leap achieved in the late stages of human evolution [6–8]. It can be seen that language abilities and cognitive abilities overlap at both the basic and higher levels, and at certain specific points they are even difficult to distinguish from each other [9–11]. This aligns well with the current state of artificial intelligence: intelligence need not depend on existing language, yet relying on advanced language abilities seems to drive and support the implementation of higher-order cognitive capacities. Nevertheless, one vaguely senses that fundamental research on the relationship between language and cognition may see new major breakthroughs in the future. Exploring the relationship between language and cognition from a genomic perspective remains a fundamental starting point.

Advanced language and cognitive abilities are key traits that distinguish humans from other primates, and their genetic basis has long been a focal point of research in evolutionary biology and neuroscience. Genome-wide association studies (GWAS) and comparative genomics have identified multiple candidate genes associated with language disorders, reading ability, intelligence, and psychiatric conditions [12–15]. For instance, *FOXP2* is considered the first discovered “language gene,” with its mutations leading to severe speech apraxia [16]; *CNTNAP2* has been linked to Specific Language Impairment (SLI) and autism spectrum disorder [17]; and *ARHGAP11B* is thought to be closely related to human cerebral cortical expansion and cognitive evolution [18]. In addition, genes such as *ASPM* play important roles in neurodevelopment and cognitive function [19–20]. These findings suggest that the molecular underpinnings of language and cognitive abilities involve multiple genes and their interaction networks, and that these genes may have experienced different selective pressures during the process of species divergence.

Against the above background, this study raises the following scientific question: during the evolution of SNVs in cognitive genes, do language-related genes and cognition-related genes exhibit significantly different intensities of change at several key evolutionary moments (i.e., the inflection points or derivative peak positions of similarity curves)? Specifically, do language genes appear more frequently than cognitive genes in mutation peak regions, and do they have higher difference scores? To address these questions, we employed nine distance calculation methods to compare the base strings at 13,415 SNV loci across the whole genomes of 413 animal samples from multiple taxa. Using the modern human sample pp6 (which is actually *Homo_sapiens.GRCh38*) as a reference, we calculated the genetic distance for each sample and converted it to a similarity in the range [0, 1] using 1/(1+distance). Scatter-plot and line graphs were drawn using raw data (without smoothing). For each method, samples were independently sorted so that the similarity curves were naturally smooth, avoiding artificial spikes. By means of differential derivation, we located the mutation points of evolutionary rate (taking the top 10 for each method). Combined with functional annotations of 33 known language/cognition genes, we quantified the occurrence frequency and difference scores of the two gene categories in the peak sample windows (all calculations were performed using R code).

## Methods

### Data sources and preprocessing

The input data include meta-information of 413 samples (Supplemental Table 1: *meta_data.xlsx*) and 13,415 SNV genotypes (Supplemental Table 2: *snv_data.xlsx*). *meta_data.xlsx* contains 7 columns: sample (sample ID), taxon, country, region, age (BP) (years before present), sample_details (detailed description of the sample), and accession (public genome sequence database source and accession number). The samples cover 11 taxonomic groups, including: Amphibian: 6 samples; Artiodactyla: 40 samples; Birds: 50 samples; Cetacea: 34 samples; Fish: 92 samples; Human: 34 samples; Laurasiatherians: 38 samples; Others: 23 samples; Primates: 33 samples; Reptiles: 34 samples; Rodents: 29 samples. These include many ancient human samples (with complete chronological information) and living fossil animals (such as coelacanth, lungfish, tuatara, etc.), spanning from fish to humans, thus providing rich material for studying cross-species evolution of cognitive genes. Except for human samples and living fossil species (e.g., lungfish, coelacanth, hagfish, horseshoe crab, *Nautilus pompilius*, pig-nosed turtle, platypus, etc.), all other samples are representative species, with no multiple samples from the same species (which would cause redundancy in similarity). The bases were encoded numerically (A=1, T=2, C=3, G=4, missing=0), and loci with all zero values were removed, retaining a total of 13,415 effective SNV loci. These SNV loci were obtained as follows: using a custom search software written in Python/C++, we searched the whole-genome sequence files (in FASTA or FASTQ formats) of each sample for the sequences of the 13,415 SNV loci from the 33 human language/cognition genes. Each SNV locus sequence consisted of the SNV base plus equal-length flanking bases on both sides, with a total length of 27 bases. We required a 100% match for the 26 flanking bases, which allowed us to obtain the base data at each SNV locus in the sample genome.

### Similarity calculation and ranking

Nine distance metrics were used: Bray–Curtis, Cosine, Pearson, Spearman, Hamming, Jaccard, Matching, Kulczynski, and Gower [21–30]. For each sample, the distance to the reference sample pp6 was calculated separately. To place the similarities from all methods into a uniformly comparable interval of (0,1], we applied the transformation *similarity* = 1/(1 + *distance*). This transformation preserves the monotonic property that smaller distances correspond to larger similarities, while ensuring that all similarity values fall within (0,1]. For each method, samples were independently sorted in ascending order of similarity (i.e., descending order of distance) along the horizontal axis, with similarity plotted on the vertical axis, so that the resulting curves are monotonically increasing.

For each method, we first compute a raw distance (d) or raw similarity (s_raw).To bring all values into a comparable [0,1] range, we apply: similarity = 1 / (1 + d) where d is the raw distance (or 1 - s_raw for methods that natively produce a similarity). Below are the specific calculations for each method:

- Bray-Curtis: raw distance = sum(|x_i - y_i|) / sum(x_i + y_i + epsilon) similarity = 1 / (1 + raw_distance)
- Cosine: raw similarity = dot_product / (||x|| * ||y|| + epsilon) raw distance = 1 - raw_similarity similarity = 1 / (1 + raw_distance) = 1 / (2 - raw_similarity)
- Pearson: raw correlation r = Pearson correlation between x and y raw distance = 1 - r similarity = 1 / (1 + raw_distance) = 1 / (2 - r)
- Spearman: raw correlation rho = Spearman rank correlation raw distance = 1 - rho similarity = 1 / (1 + raw_distance) = 1 / (2 - rho)
- Hamming: raw distance = proportion of mismatched positions similarity = 1 / (1 + raw_distance)
- Jaccard: raw distance = 1 - (intersection / union) for binary data similarity = 1 / (1 + raw_distance)
- Matching: raw distance = 1 - (sum(x == y) / length) similarity = 1 / (1 + raw_distance)
- Kulczynski: a = sum(min(x_i, y_i)), b = sum(x_i), c = sum(y_i) raw distance = 1 - 0.5*(a/b + a/c) similarity = 1 / (1 + raw_distance)
- Gower: For each variable, range = max - min, scaled_diff = |x_i - y_i| / range raw distance = mean(scaled_diff) similarity = 1 / (1 + raw_distance)

### Derivative and peak detection

Given the specific nature of the similarity curves in this study, derivatives were calculated from adjacent similarity differences, excluding the first and last 5 samples on each side to avoid boundary effects. The top 10 points with the largest absolute derivative values were selected as mutation peaks of evolutionary rate.

### SNV association and gene annotation

For each peak, we took the peak sample together with 8 samples before and after it (adjustable) as a window (17 samples in total). We calculated the difference score (number of mismatched samples) at each locus between the peak sample and the other samples in the window, selected the top 600 loci (adjustable) with the highest scores, and mapped them to genes.

### Definition of quantitative metrics

The “occurrence frequency” was defined as the proportion of differential SNVs belonging to language (or cognition) genes among all differential SNVs within a given peak window. The “difference score proportion” was defined as the ratio of the sum of difference scores of a particular gene category to the total difference score.

## Results

### Similarity curves and derivative peaks

The samples in Figure 1 cover a wide range, with a relatively large sample size, including representative samples from all major animal groups, and each sample contains 13,415 effective SNV loci. This means that the basic shape of the similarity curve in Figure 1 has been largely determined, and adding more samples is unlikely to significantly alter its overall profile. In fact, over a recent period, the authors have applied nearly 30 different methods for calculating similarity based on SNV base strings, and the vast majority produced contour curves highly similar to that in Figure 1; this paper reports only nine of those methods. Therefore, using this curve to model the dynamic evolutionary process of cognitive gene polymorphism patterns is practically reliable. Given the large number of samples, the density of points in the scatter-plot curve is sufficiently high, so that the curve appears very similar to a smoothed version, and the steep slopes on the curve can reasonably reflect transitions in the pattern of SNV polymorphism (note that this reflects changes in multi-locus patterns, not mutations at individual loci). This is also an important reason why derivatives calculated from adjacent similarity differences can serve as a reliable approximation in this study.

**Figure 1.**
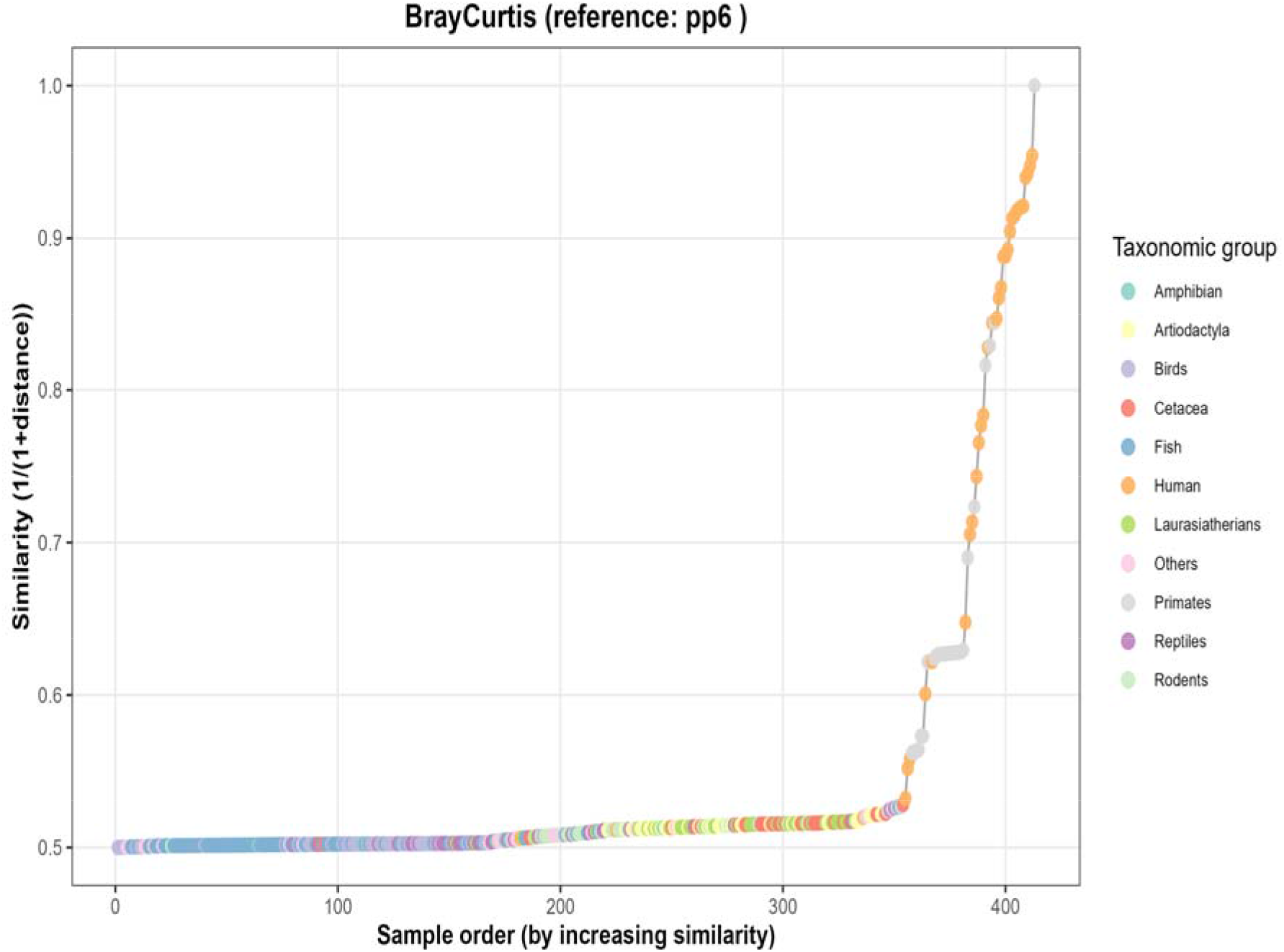
Raw scatter plot with connecting lines for Bray–Curtis similarity. Horizontal axis: sample indices sorted in ascending order of similarity; vertical axis: similarity (1/(1+distance)). Each circle represents one sample, with colors indicating taxonomic groups. The grey line connects adjacent samples, showing the trend of similarity changes along the sorting order. Because the samples are independently sorted, the curve is monotonically decreasing and free of artificial spikes.

Due to the relatively large number of samples, the similarity curves calculated by multiple methods (including other series of methods beyond the nine presented here, data not shown) consistently reveal small steps of early mutations, or more precisely, three notably rapid upward jumps in similarity within the horizontal-axis range of approximately 350–385. After these three jumps, the evolutionary rate of cognitive/language genes remains generally high, albeit with several apparent plateau-adjustment periods in between (Figures 1–4). These small steps were checked and are not due to multiple samples from the same species (e.g., living fossil species generally have 2–3 samples per species in the dataset).

When the derivative curves of the nine methods are displayed side-by-side (Figure 5), one can observe at which positions large mutations (peaks) occur across the methods. Most methods show dense peaks between indices 350 and 400, indicating that this region is a critical interval for changes in evolutionary rate under multiple distance metrics. A closer look reveals that the positions around 355, 363, and 382 are where the evolutionary rate of cognitive/language genes first shows significant changes, corresponding to the two small plateaus on the similarity curve and the three steep rises to the left of those plateaus.

**Figure 2.**
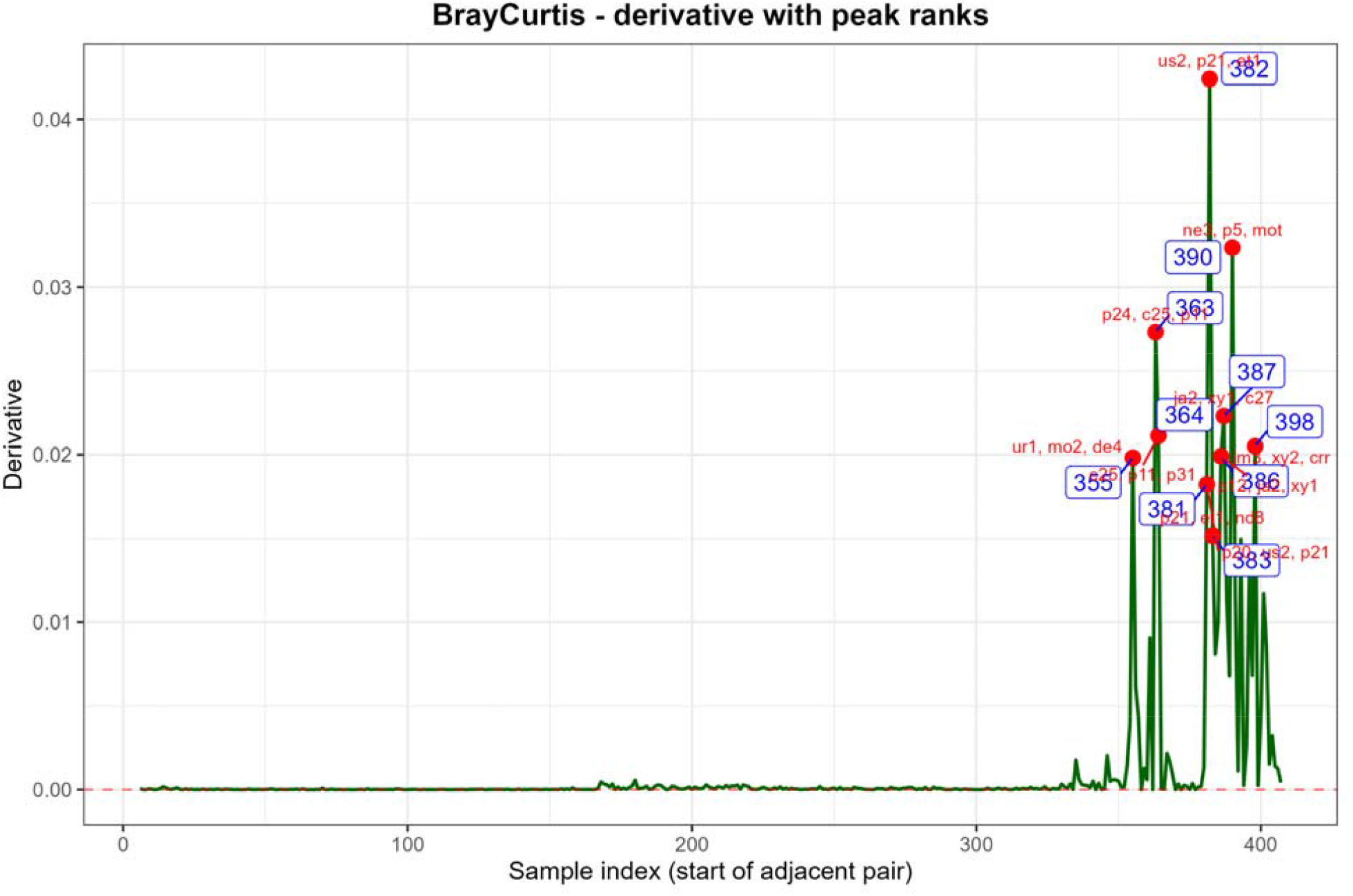
First-derivative curve of similarity. Horizontal axis: starting index of adjacent sample pairs (excluding the first and last 5 on each side); vertical axis: difference values of similarity. The red dashed line is the zero reference line, and red circles mark the ten peak positions with the largest absolute derivative values.The names and data sources of all peak samples are provided in the meta_data table.

**Figure 3.**
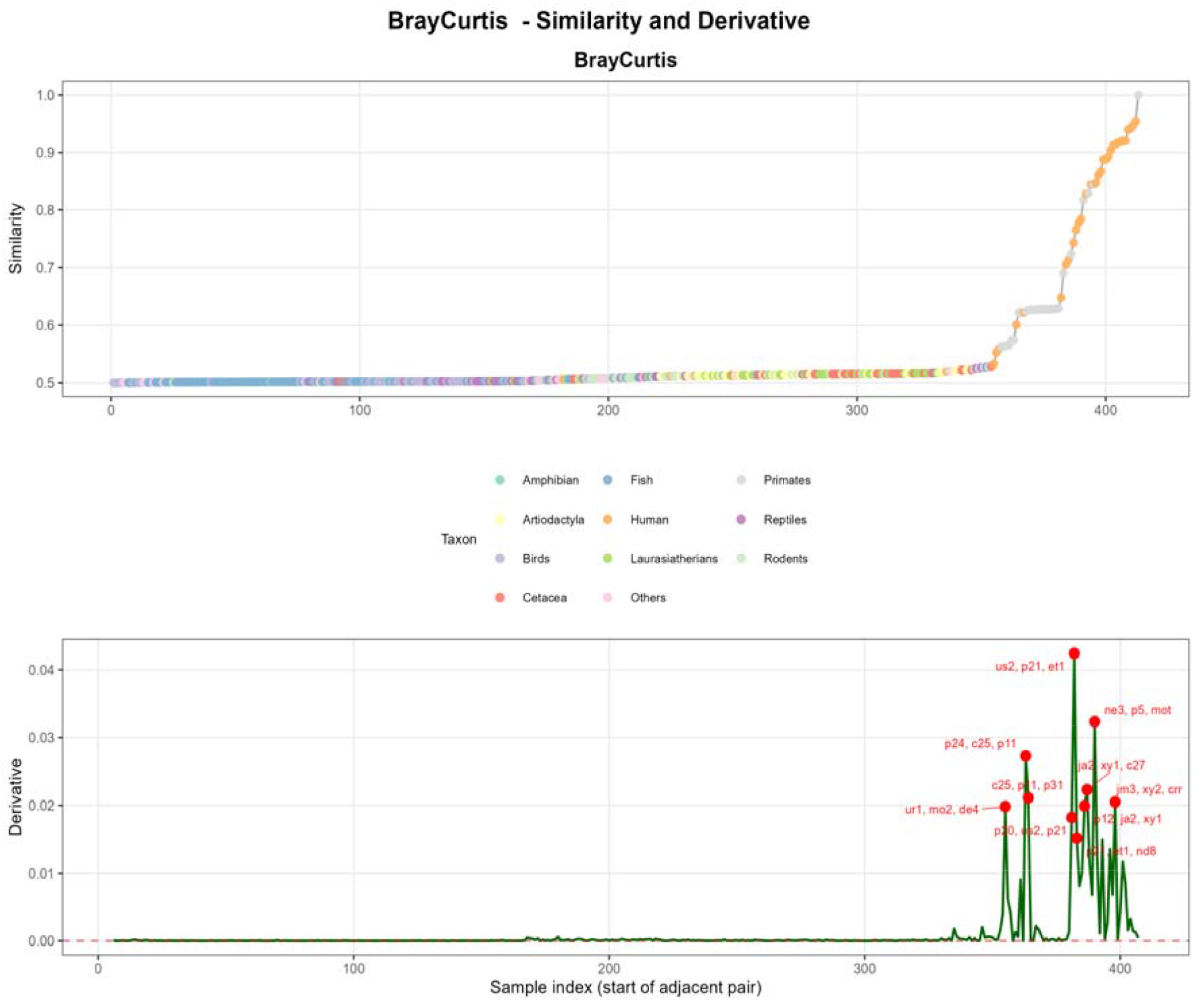
Composite plot of the similarity curve and the derivative curve. The upper panel shows the similarity scatter-plot with connecting lines, and the lower panel shows the derivative curve. Red points mark the top 10 peak positions, with corresponding sample names labelled. This figure facilitates the correspondence between steep slopes and small plateaus on the similarity curve and the peaks on the derivative curve.

**Figure 4.**
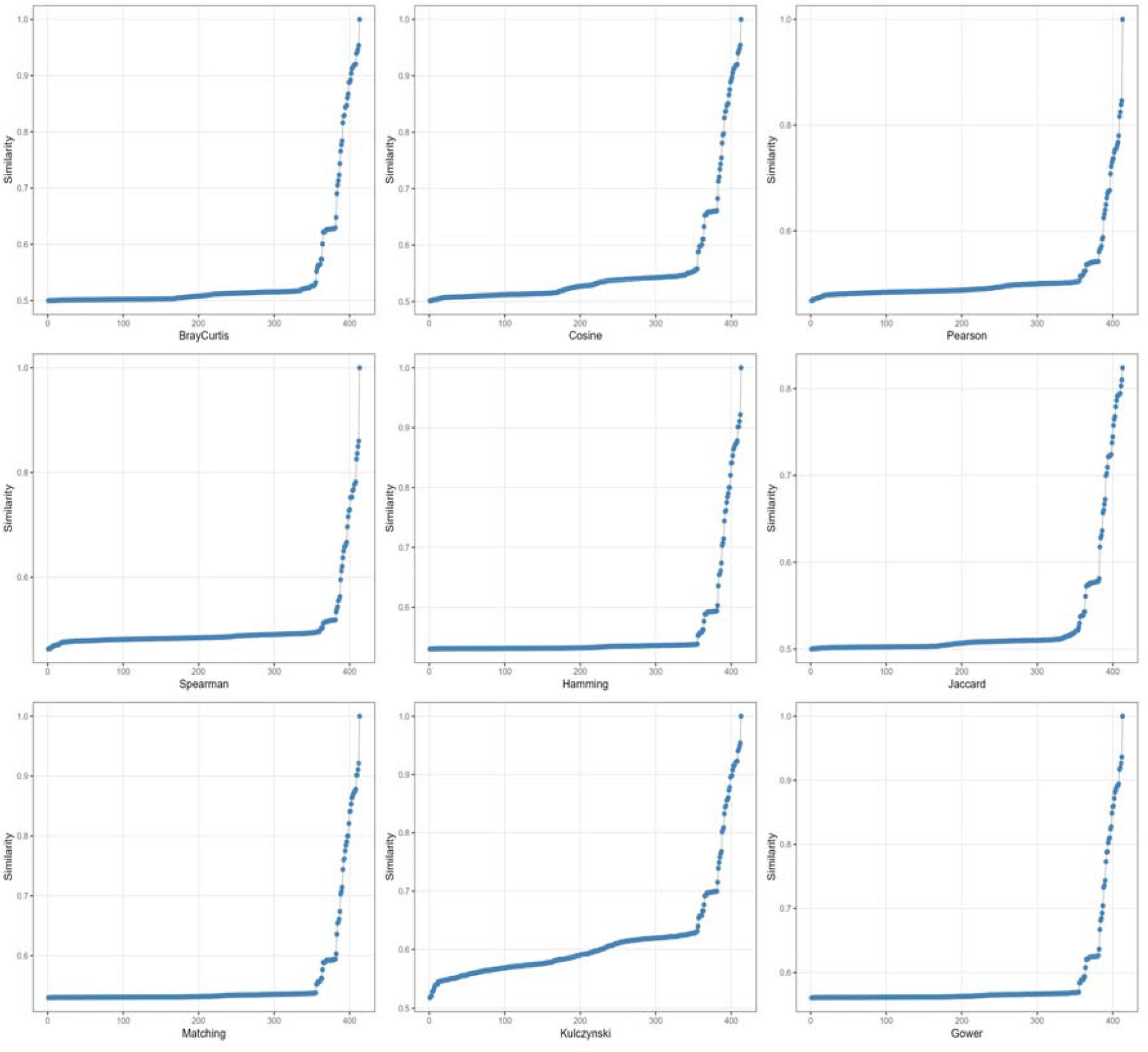
The similarity curves from the nine methods are arranged together for easy horizontal comparison of the influence of different methods on the similarity ranking. For each method, samples are independently sorted according to their similarity values; all curves are monotonically increasing, although the rates of ascent and the curve shapes differ slightly.

**Figure 5.**
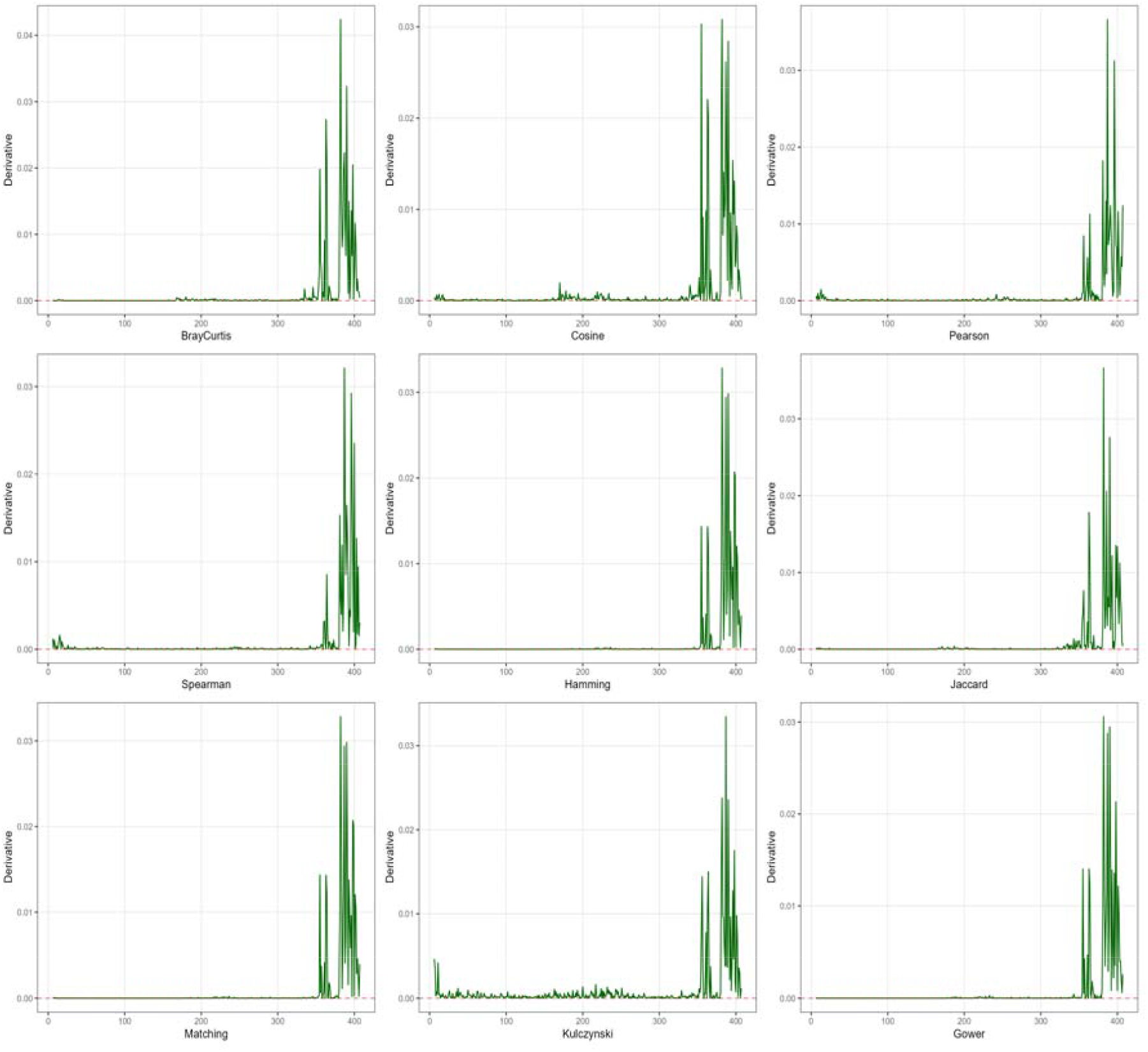
The nine derivative curves are shown together. It can be seen that the nine similarity calculation methods differ in their sensitivity to detecting the first three steep slopes and two small plateaus that appear on the similarity curves (corresponding approximately to the horizontal-axis positions of 355, 363, and 382).

**Figure 6.**
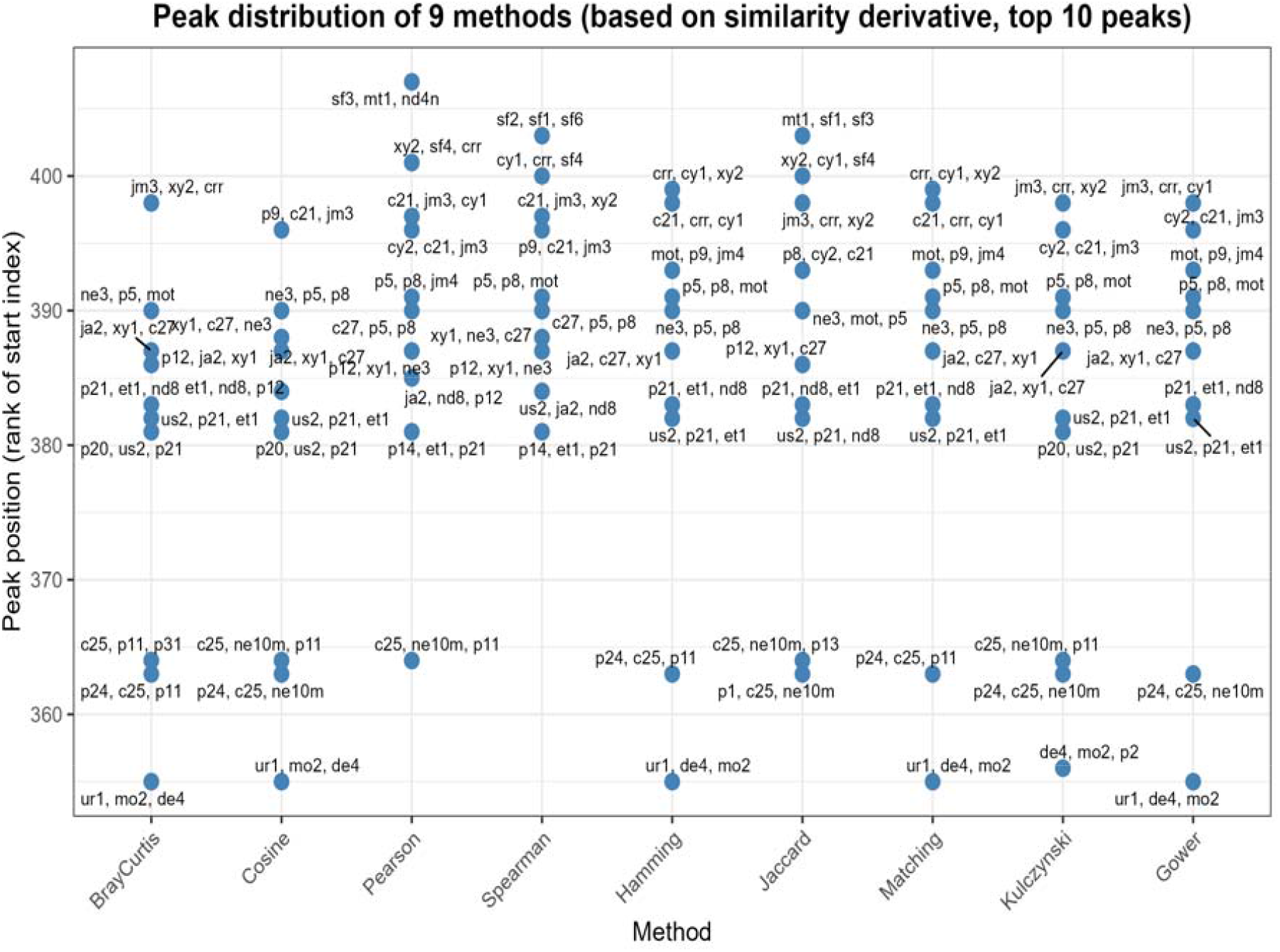
Distribution of peak positions detected by the nine distance methods. Horizontal axis: method; vertical axis: starting sample index corresponding to each peak. Each method has 10 peak points. The peak position and the names of the two adjacent samples are annotated beside each point. The names and data sources of all peak samples are given in the meta_data table.

The boxplots in Figure 7 show the peak derivatives detected by each method. In principle, since for all methods the similarity curves are plotted with samples sorted in ascending order of similarity along the horizontal axis, and the derivative values (as shown in Figures 2 and 5) are all positive, larger vertical-axis values in Figure 7 should correspond to greater magnitudes of mutation. Clearly, the two samples p21 (*Nomascus leucogenys*, Ensembl) and xy1 (Xingyi_LN, Yunnan, PRJCA015361) [31] have the largest peak derivatives; it is noteworthy that two primate samples (p21 and p5, *Gorilla gorilla*, Ensembl) are among those with the largest derivative peaks. Several other samples with very large derivative peaks are c21 (LJM3, China, PRJEB36297), c27 (HJTM115, China, PRJEB36297), and mo2 (Iberomaurusian, Morocco, PRJNA422662) [32–33]. It is worth noting that the three peak samples with the largest mutation magnitudes (xy1, c21, c27) are all from East Asia. Whether this phenomenon is related to the fact that East Asian populations have the largest average brain size globally merits further attention.

**Figure 7.**
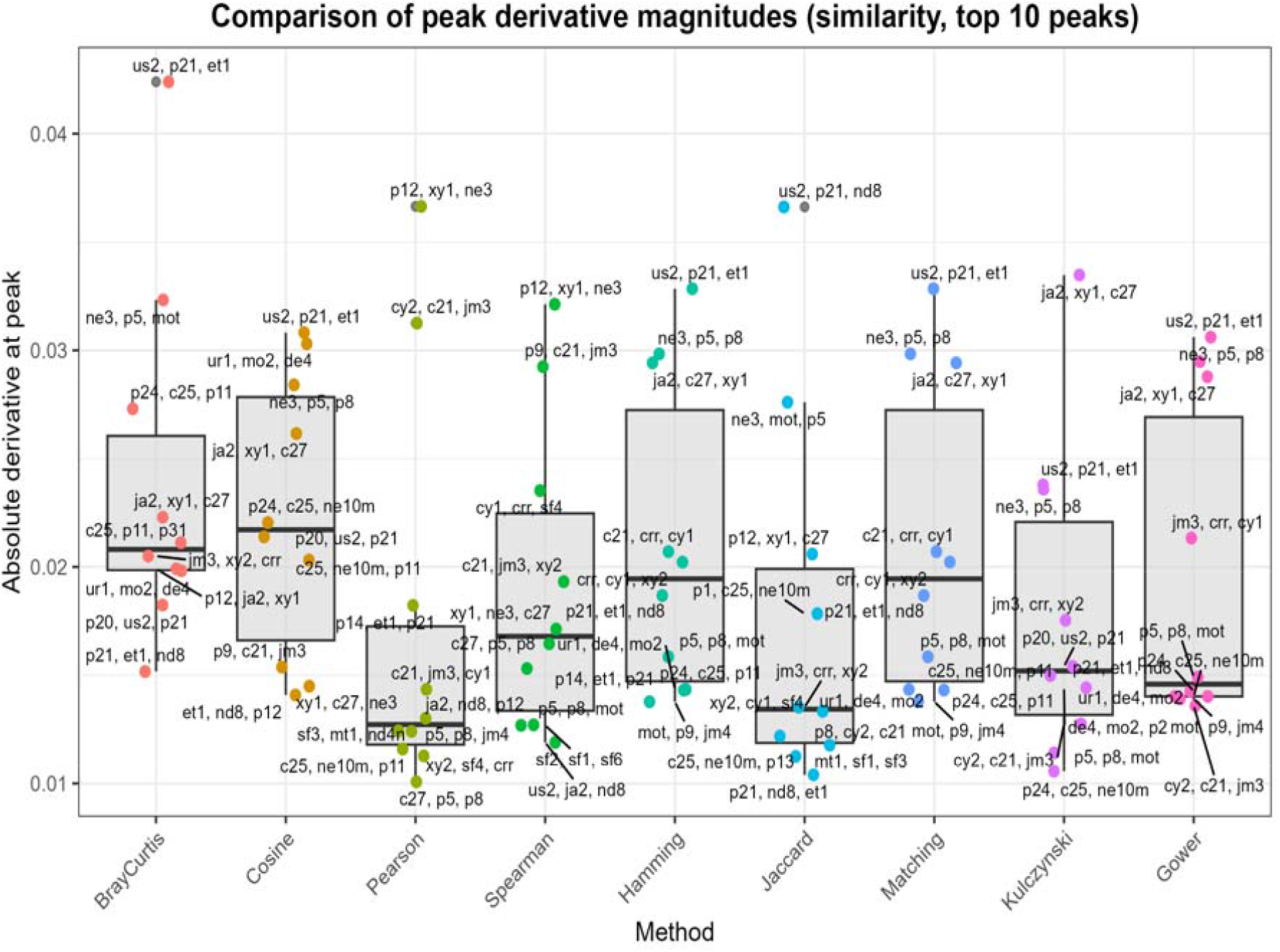
Boxplots of the absolute derivative values at the peaks for each method. Horizontal axis: method; vertical axis: absolute derivative value at the peak. The box represents the interquartile range of the 10 peaks, with overlaid scatter points showing the exact values for each peak; sample names are annotated beside the points. The Gower and Hamming methods have the widest boxes, suggesting that they may be more sensitive to changes in similarity. The names and data sources of all peak samples are given in the meta_data table.

The shape of the boxplots in Figure 8 reflects that the similarity data are partitioned into two parts: one part consists of samples with very small values, and the other part consists of samples with rapidly increasing similarity values. These two groups of data are also clearly visible in Figures 1, 3, and 4. This indicates that the polymorphism of cognitive/language genes in animal taxa began to accelerate suddenly at a certain stage of evolution, and this acceleration appears to occur mostly in primates and ancient human samples.

**Figure 8.**
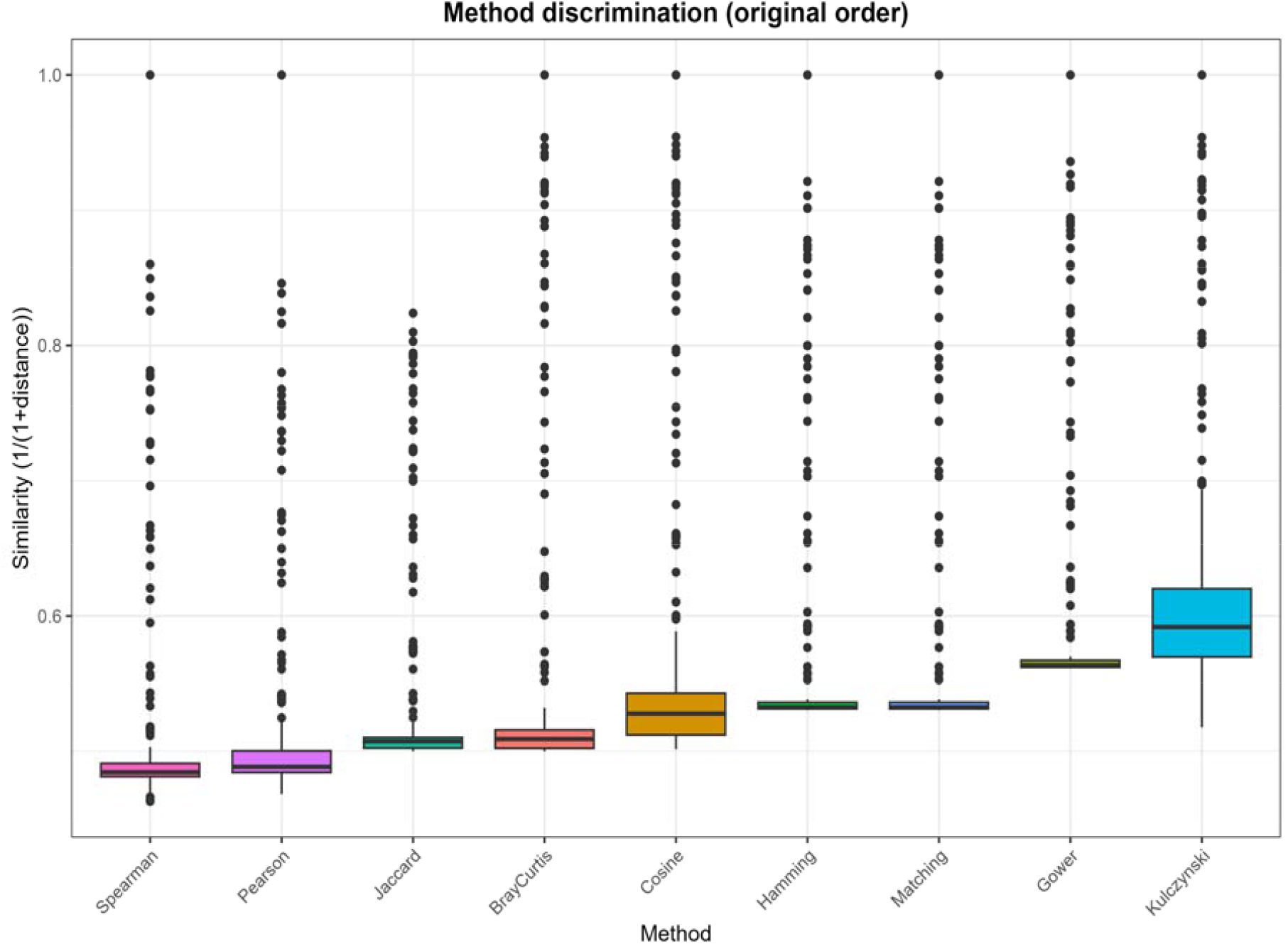
Boxplots of similarity across all samples for the nine methods. Horizontal axis: method names (sorted by median); vertical axis: similarity (1/(1+distance)). The box represents the median and interquartile range, with outliers shown as points. A wider box indicates greater discriminative power of the method for sample differences. The Kulczynski and Cosine methods have the widest boxes, suggesting that they may be more sensitive to SNV differences.

### Association between gene function and peak SNVs

We tested N values of 300, 500, 600, 800, 1,000, 1,200, 1,500, and 1,800, and found that only when N is at least 600 can the unique SNV loci of the peak samples be fully extracted from the flanking samples. At N=600, among the differential SNVs in all peak regions, the total difference score for language-related genes was 436,843 (accounting for 54.6% of the total difference score), and for cognition-related genes it was 468,324 (58.6%). The sum exceeds 100% because some genes are associated with both language and cognition functions (i.e., overlap). The average difference score per gene was 22,991.74 for language genes and 26,018 for cognition genes. The average variation intensity per locus was 267.55 for language genes and 295.53 for cognition genes. This indicates that the occurrence probabilities of unique SNVs of language genes and cognition genes at peak positions are very similar, with cognition genes showing slightly higher occurrence frequencies and per-locus variation intensities (see Table 1 for the top 6 difference scores of language and cognition genes).

**Table 1.**
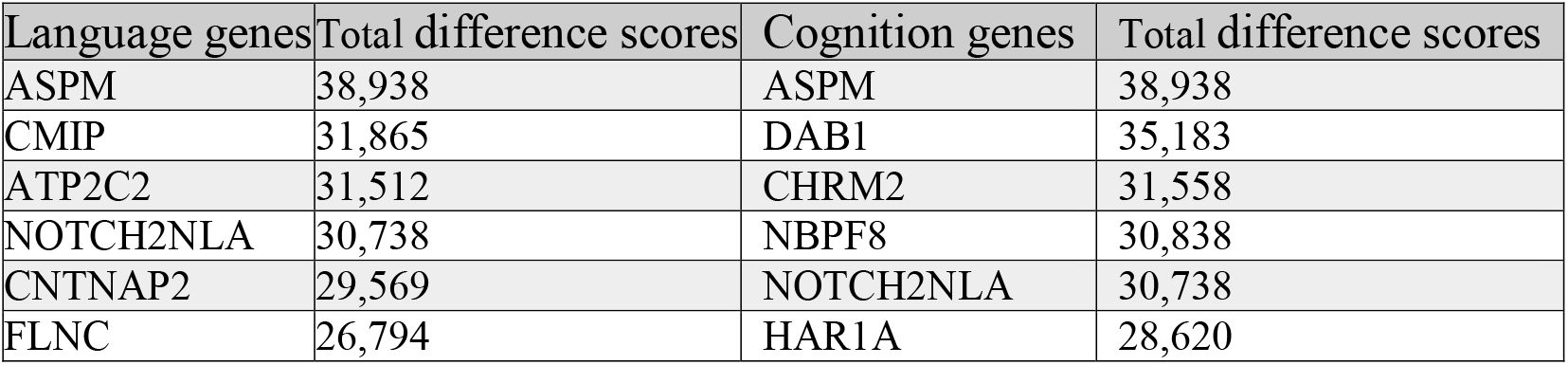
The top 6 difference scores of language and cognition genes.

### Analysis of SNV differences between peaks and left-side samples

In this analysis, we took the sample immediately to the right of each peak (Start_Index + 1) as the “peak sample,” because the derivative peak reflects the mutation from the left-side sample to the right-side sample. We then compared this peak sample, locus by locus, with the six samples on its left, recording those loci where the peak sample differed from all left-side samples, and then computed the intersection of such differing loci for the peaks within each corresponding region. In the 355–356 region, we found one intersection locus, corresponding to a language-related gene (*SRGAP2*); in the 381–382 region, there were two intersection loci, corresponding to language-related genes (*NFXL1* and *SRGAP2*). In the regions 363–364, 383– 385, 386–388, 390–393, 396–399, and ≥400, no intersection loci were found.

However, if we compare the peak sample only with its immediate left neighbor in terms of SNV differences, the results may differ. Based on the results from the nine methods as shown in Figure 2, we obtained three groups of peaks:

- Group 1: Hamming (355), Kulczynski (356), Matching (355), Bray–Curtis (355), Cosine (355), Gower (355);
- Group 2: Hamming (363), Jaccard (363), Kulczynski (363, 364), Matching (363), Pearson (364), Bray–Curtis (363), Cosine (363, 364), Gower (363);
- Group 3: Hamming (382), Jaccard (382), Kulczynski (381, 382), Matching (382, 383), Pearson (381), Spearman (381), Bray–Curtis (381, 382), Cosine (381, 382), Gower (382, 383).

These three groups of peaks correspond to the horizontal-axis positions 355, 363, and 382 on the similarity curves in Figure 1 (across the nine panels), which are the locations where the evolutionary rate of cognitive/language genes first shows significant changes. For each group of peaks, we performed the following: for all peak samples (Start_Index + 1) in that group, we took the intersection of their entire set of 13,415 SNVs to obtain intersection set 1; similarly, we took the intersection of the SNVs from all the immediate left-neighbor samples (Start_Index) in that group to obtain intersection set 2. We then identified the SNV loci that are present in intersection set 1 but absent in intersection set 2, and mapped them to their corresponding genes. Using this approach, we found that the intersecting differential loci in Group 3 involved three language-related genes (*NFXL1, SRGAP2*, and *SRGAP2C*). This result is consistent with that obtained from the previous method (comparing the peak sample with its six left-side samples). Of course, this finding does not prove that the substantial similarity jump corresponding to the derivative peaks is caused by mutations in language genes; nevertheless, it does suggest, to a certain extent, that SNV mutations in language genes may have played a distinctive role.

## Discussion

In this study, nine similarity (distance) calculation methods were used to plot similarity curves based on the base sequences at 13,415 SNV loci from 413 animal samples across various taxa, and a series of analyses were subsequently carried out. The specific ordering along the horizontal axis of the similarity curves makes the curves monotonically increasing, with several steep slopes appearing. Due to the relatively large sample size, the density of points on the curves is high, so that the scatterplots appear very similar to smoothed curves. Under these circumstances, using the differences between adjacent samples as a proxy for derivatives can indeed reflect the intensity of changes in the multi-locus SNV polymorphism patterns under this sample condition. If the sample size were to increase further in the future, these steep slopes might disappear. However, it should be noted that there are some small plateaus on the similarity curves [these plateaus are not artifacts caused by redundant sampling of highly similar samples or samples from the same species]. This implies that even if the sample size were increased a hundredfold, these small plateaus would not disappear. On either side of these small plateaus, steep slopes will inevitably arise; in other words, mutations in the SNV polymorphism patterns of language/cognition genes are inherently present. These steep slopes may conceal several key secrets of the evolution of cognitive functions.

When the full set of SNVs from language genes and the full set of SNVs from cognition genes were separately subjected to similar analyses on the 413 samples, it was found that the similarity curves for both categories of genes were highly consistent with those in Figures 1 and 3; the positions of steep slopes and small plateaus on the curves were also highly consistent (data not shown). For language genes alone, the six language genes with the highest proportion of difference scores at peak positions (ranked by total difference score) were: *ASPM* (47,777); *NOTCH2NLA* (44,322); *ATP2C2* (41,851); *CMIP* (40,404); *SRGAP2* (37,842); and *FLNC*

(37,405). For cognition genes alone, the six cognition genes with the highest proportion of difference scores at peak positions (ranked by total difference score) were: *ASPM* (51,133); *DAB1* (44,603); *CHRM2* (40,774); *ARHGAP11B* (40,087); *NBPF8* (38,644); and *HAR1A*

(37,922). These results are largely consistent with those in Table 1.

The original intention of this study was to explore, at specific positions on the similarity curves, whether quantitative differences between language genes and cognition genes could be detected. Indeed, we found that at the three steep slopes of the similarity curves, all methods consistently identified SNV mutations in three language genes (*NFXL1, SRGAP2*, and *SRGAP2C*). When we separately analyzed the SNVs of language genes and those of cognition genes through the entire pipeline, we found that at the third steep slope peak, both sets included mutations in the language gene *SRGAP2C*, and both also included mutations in the cognition genes *DAB1* and *NRXN1*. These are preliminary results, and they only tentatively suggest that certain specific SNV loci of the two gene categories exhibit definite frequency differences at the existing steep-slope positions.

This also validates, from another angle, the results already obtained in this study: the SNV polymorphisms of the two gene categories have undergone changes of similar magnitude during evolution, so that the degree and rate of similarity change are relatively consistent. Although cognition-related genes show slightly higher occurrence frequencies in the peak windows and slightly higher average difference scores per gene than language genes, the evolutionary trajectories of the two gene categories appear to have maintained a comparable pace, with differences only at a small number of SNV loci at the peak positions. Whether these minor differences have a major impact on the evolutionary process of cognitive abilities remains unknown at present.

In this study, the top-ranked genes by total difference score included *ASPM, DAB1, ATP2C2, CHRM2, CMIP*, and *CNTNAP2*. These genes are known to play critical roles in cognitive and language evolution. For example, *ASPM* is associated with cerebral cortical size and microcephaly [38]; *DAB1* is related to neuronal migration and cognitive function [39]; *ATP2C2* shows significant association with language impairment [40]; *CHRM2* plays an important role in cognitive function [41]; *CMIP* has been identified as a novel candidate gene for neurodevelopment and psychiatric disorders [42]; and *CNTNAP2* is a classic communication- and language-related gene [43]. The concentration of difference scores for these genes at evolutionary inflection points further supports their contribution to the evolution of cognitive and language abilities.

Across all detected peak windows, a total of three living-fossil samples appeared in at least one peak, with a combined occurrence of seven times. Among them, the most frequently observed were lu3 (West African lungfish, PRJNA813995), vk2 (Komodo dragon, PRJNA523222), and pn2 (pig-nosed turtle, PRJNA512133), appearing in 3, 3, and 1 peak windows, respectively. These living-fossil species occupy key branching positions in evolution; the mutation peaks of their SNV loci within cognitive gene regions suggest that these ancient taxa may carry ancestral polymorphisms related to neurodevelopment.

Across all peak windows, a total of 26 ancient human samples appeared, with a combined occurrence of 778 times. The five ancient human samples with the highest occurrence frequencies were c27, ne3 (Kyang-KS20, PRJEB41752), mot (Mota, PRJNA295861), ja2 (Funadomari23 female-2, PRJDB7235), and xy1, with occurrence counts of 58, 56, 54, 51, and 51, respectively.

Among the top three peaks for each method (i.e., the three evolutionary breakpoints or inflection points with the largest absolute derivative values), the ancient human samples with the highest occurrence frequencies were ne3 (occurring 9 times), et1 (occurring 7 times; Ancient Ethiopian Mota genome, PRJNA295861), and us2 (occurring 7 times; US ancient Anzick, PRJEB29074). The high frequency of these samples at the most drastic evolutionary inflection/breakpoints suggests that they may carry important variants related to the evolution of cognitive functions.

## Conclusion

Based on 413 whole-genome sequence samples, this study compared the polymorphism patterns of cognitive genes across major animal taxa at 13,415 SNV loci through similarity analysis. We found several distinct steep slopes on the similarity curves, which may correspond to important mutations in a set of SNV loci. We did not apply any smoothing to the similarity curves; instead, given that the scatter-point density was sufficiently high, we used differences between adjacent similarity values to approximate derivatives, thereby maximising the precision in locating potential key mutations. Theoretically, significant SNV changes could be hidden anywhere on the curves, and these steep slopes might simply be gaps caused by insufficient representative samples. Nevertheless, searching for the SNV pattern changes and associated genes corresponding to the existing steep slopes is methodologically meaningful (the analytical pipeline of this study can be applied to any interval on the similarity curves).

The current preliminary conclusion is that, for the high-scoring SNV loci of the two gene categories, their occurrence frequencies at the right-side positions of the several steep slopes on the similarity curves are very close, with cognition-related genes being slightly higher. However, individual SNV loci in language genes (*NFXL1, SRGAP2, SRGAP2C*) were detected at the peaks by all nine methods more often than those in cognition genes. While this suggests that certain mutations in language genes may have played a distinctive role in the overall evolution of cognitive abilities, this result remains preliminary observational data at present.

## Acknowledgments

This study was supported by a State Language Commission Research Grant (YB135-117) and National Research Center for Foreign Language Education Grant (ZGWYJYJJ10A042). The authors are indebted to Shuaiyu Zhang for helping in SNV-searching software development.

